# Loss of TDP-43 function drives cryptic circular RNAs in neurodegenerative diseases

**DOI:** 10.64898/2026.09.17.752169

**Authors:** Chengzhang Zhu, Zhiyan Zhao, Bojun Song, Yan-Ming Chen, Yu Xiao, Zhe Zhang, Yingzhi Ye, Niannian Xu, Ruijia Zhang, Yongxin Huang, Juan C. Troncoso, Chuan He, Shuying Sun

**Affiliations:** Department of Department of Physiology, Pharmacology & Therapeutics, Johns Hopkins University School of Medicine, Baltimore, MD 21205, USA; Brain Science Institute, Johns Hopkins University School of Medicine, Baltimore, MD 21205, USA; Cellular and Molecular Medicine Graduate Program, Johns Hopkins University School of Medicine, Baltimore, MD 21205, USA; Biochemistry and Molecular Biology Program, Johns Hopkins Bloomberg School of Public Health, Baltimore, MD 21205, USA; Department of Chemistry, The University of Chicago, Chicago, IL, 60637, USA; Department of Biochemistry and Molecular Biology, The University of Chicago, Chicago, IL, 60637, USA; Institute for Biophysical Dynamics, The University of Chicago, Chicago, IL, 60637, USA; Howard Hughes Medical Institute, Chicago, 60637, IL, USA; Cellular and Molecular Physiology Graduate Program, Johns Hopkins University School of Medicine, Baltimore, MD 21205, USA; Department of Pathology, Johns Hopkins University School of Medicine, Baltimore, MD 21205, USA; The Solomon H. Snyder Department of Neuroscience, Johns Hopkins University School of Medicine, Baltimore, MD 21205, USA

## Abstract

TAR DNA-binding protein 43 (TDP-43) is a key pathological hallmark of amyotrophic lateral sclerosis (ALS) and frontotemporal dementia (FTD) and a critical regulator of RNA splicing. While loss of TDP-43 induces aberrant splicing in linear transcripts, its impact on circular RNA (circRNA) biogenesis remains unexplored. Here, we show that TDP-43 depletion in human neurons induces widespread circRNA changes, especially upregulation of a distinct class of cryptic circRNAs that arise specifically upon loss of TDP-43. Some of these cryptic circRNAs incorporate cryptic exons derived from intronic sequences. Notably, these cryptic circRNAs exhibit greater stability than their corresponding linear RNA isoforms and accumulate progressively in neurons. Moreover, cryptic circRNAs are elevated in postmortem brain tissues from ALS, FTD and Alzheimer’s disease (AD) patients. These findings reveal a previously unrecognized role for TDP-43 in repressing cryptic circRNA formation and establish these circRNAs as stable molecular signatures of TDP-43 dysfunction.

## INTRODUCTION

Amyotrophic lateral sclerosis (ALS) and frontotemporal dementia (FTD) are neurodegenerative disorders that share common genetic, clinical, and pathological features^1–3^. A defining hallmark of both diseases is the proteinopathy of TAR DNA-binding protein 43 kDa (TDP-43), characterized by its nuclear depletion and cytoplasmic mislocalization in affected neurons and glia^4, 5^. Beyond ALS and FTD, TDP-43 pathology is observed across a broad range of neurological disorders, including Alzheimer’s disease (AD)^6–8^, Parkinson’s disease^9, 10^, Huntington’s disease^11^, limbic-predominant age-related TDP-43 encephalopathy (LATE)^12^, and multiple sclerosis^13^.

Emerging evidence indicates that nuclear depletion of TDP-43 frequently precedes overt cytoplasmic aggregation during disease progression^14, 15^. In the nucleus, TDP-43 functions as a critical regulator of pre-mRNA splicing, and its loss leads to aberrant mis-splicing events such as cryptic exon (CE) inclusion, intron retention, and exon skipping^16–21^. Beyond splicing, TDP-43 also regulates multiple aspects of RNA metabolism, such as alternative polyadenylation^22–25^ and mRNA stability^26, 27^. Besides mRNA, increasing evidence also highlights novel roles of TDP-43 in non-coding RNA regulation, such as microRNAs^28–30^ and retrotransposon RNAs^31–34^. Together, these RNA processing defects resulting from TDP-43 loss-of-function (LOF) contribute to neuronal dysfunction and degeneration, supporting TDP-43 LOF as a convergent molecular mechanism across neurodegenerative conditions^17, 20, 35–37^.

Circular RNAs (circRNAs) are covalently closed, single-stranded RNAs that are generated through spliceosome-mediated back-splicing^38, 39^. Because they lack free 3′ and 5′ ends, circRNAs are resistant to exonuclease-mediated degradation and are relatively refractory to RNA decay pathways in the cell^39–42^. CircRNAs are found to be mostly enriched in the nervous system, where they have been implicated in neuronal gene expression, development, and synaptic function^43, 44^. Emerging evidence suggests that circRNA dysregulation occurs in neurodegenerative diseases, such as Alzheimer’s disease^45, 46^ and Parkinson’s disease^47, 48^. However, the underlying molecular mechanisms remain poorly understood. CircRNA biogenesis is tightly coupled to canonical splicing regulation and can be promoted by RNA-binding proteins (RBPs) that bind to flanking intronic regions to facilitate back-splicing through RNA looping^42, 49^. Given that TDP-43 exhibits strong binding enrichment in introns, we examined whether loss of TDP-43 influences circRNA biogenesis.

In this work, we identify TDP-43 as a regulator of back-splicing and circRNA biogenesis, uncover a previously unrecognized class of cryptic circRNAs induced by loss of TDP-43, and demonstrate that these TDP-43–repressed circRNAs are upregulated in human neurodegenerative disease tissues. These circRNAs may persist as stable molecular consequences of splicing dysregulation, even when the corresponding linear transcripts are rapidly turned over, thereby potentially contributing to disease pathogenesis and serving as candidate biomarkers.

## RESULTS

### Loss of TDP-43 induces cryptic circRNAs in i^3^Neurons

To examine whether loss of TDP-43 affects circRNA expression in neurons, we differentiated i^3^Neurons^50^ carrying the CRISPRi machinery together with constitutive expression of an sgRNA targeting TDP-43. Efficient reduction of TDP-43 protein was confirmed by immunoblotting after 14 days of differentiation (Fig. 1a). We enriched circRNAs prior to RNA sequencing by removing poly(A)+ mRNAs, followed by RNase R treatment and rRNA depletion (see Methods). In parallel, we performed RNA-seq on the poly(A)-enriched fraction to quantify mRNA expression.

**Figure 1.**
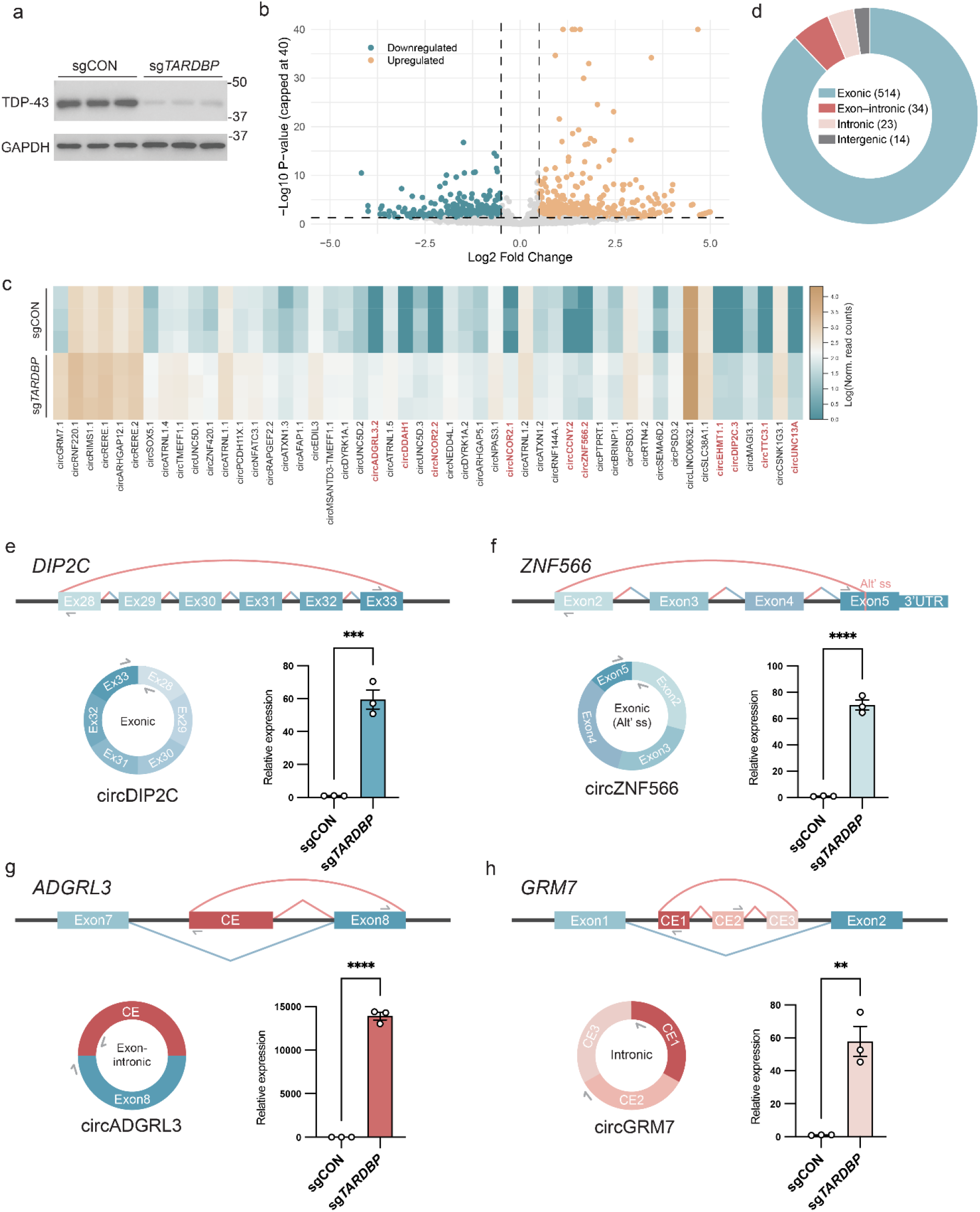
Loss of TDP-43 drives cryptic circRNA biogenesis. **a,** Western blot showing TDP-43 knockdown by sgRNA in i^3^Neurons at day 14 of differentiation. GAPDH served as a loading control. N=3 biological replicates. **b,** Volcano plot showing circRNA expression changes following TDP-43 depletion in the circRNA-enriched RNA-seq library. For visualization, the -log_10_(p-value) was capped at 40, and values above 40 are plotted at the upper boundary. A total of 7,689 circRNAs were detected, including 585 upregulated and 379 downregulated circRNAs after filtering by |LFC|>=0.5, p < 0.05. **c,** Heatmap showing log-transformed normalized read counts of the top 50 upregulated circRNAs ranked by adjusted *P* value. CircRNA isoform numbers were assigned from high to low abundance for circRNAs originating from the same host gene. Red labels indicate de novo circRNAs, defined as upregulated circRNAs with normalized read counts of 0 in at least 2 of 3 sgCON replicates. **d,** Pie chart showing categories of upregulated circRNAs based on exon–intron composition. **e–h,** Schematic representations of different types of circRNA biogenesis induced by loss of TDP-43 and an example in each category, together with RT–qPCR validation using total RNA from i^3^Neurons without circRNA enrichment. Donut graphs show the exon–intron composition of each circRNA. Canonical exons are shown in blue and CEs in red. Rounded arcs indicate circRNA backsplicing junctions and angular arcs indicate linear splicing junctions. Blue angular arcs denote the junctions used in canonical linear RNA isoforms, red arcs denote cryptic circRNA-associated junctions, and bi-colored blue–red arcs indicate splice junctions shared between canonical and circular transcripts. Arrows indicate divergent primers used for RT–qPCR amplification of circRNA back-splice junctions. Data are mean ± SEM from three biological replicates. ** P < 0.01, *** P < 0.001, **** P < 0.0001 by two-tailed unpaired *t* test.

RNA-seq analysis revealed widespread alterations in circRNA abundance following TDP-43 depletion (Fig. 1b, Supplementary Table S1). These changes showed only weak association with expression of the corresponding linear transcripts (Supplementary Fig. 1a), suggesting that circRNA dysregulation occurs largely independent of host gene transcription. Because multiple circRNA isoforms can arise from the same host gene, we assigned isoform numbers to each circRNA based on relative abundance. However, because isoform-specific analyses are not a major focus of this study, we refer to circRNAs by their host gene names throughout the remainder of the manuscript, with corresponding isoform numbers provided in Supplementary Table S1 and indicated in the main text only when necessary. Notably, we observed a more pronounced upregulation of circRNAs compared to downregulation upon TDP-43 loss (Fig.1b, c). Of particular interest, a subset of the upregulated circRNAs was nearly undetectable in control neurons and appeared predominantly de novo following TDP-43 depletion (Fig.1c, highlighted in red; Supplementary Table S2).

Next, we examined the upregulated circRNAs based on their transcriptomic features to investigate the mechanism underlying their upregulation, and validated selected candidates using divergent primers spanning the back-splice junctions. The majority of upregulated back-splicing junctions occurred between canonical exons (Fig.1d, Supplementary Table S1). Many were present in control neurons and dramatically increased upon TDP-43 loss, such as circSOX5.1, circZNF420, circTMEFF1, circRIMS1 (Supplementary Fig.1b). Some were specific to TDP-43 knockdown neurons. circDIP2C consists of six annotated exons, with the back-splicing junction connecting exon 33 to exon 28 that emerged only upon TDP-43 depletion (Fig.1e). Other validated examples include circUNC13A, circNCOR2, circDDAH1 and circCCNY (Supplementary Fig. 1b). We also observed ones with back-splicing junctions that utilized alternative splice sites within exons. circZNF566 and circSELENOT are composed of annotated exons, but their back-splicing reflects an alternative splice site usage inside exon 5 and exon 6 (Fig.1f, Supplementary Fig. 1c).

Intriguingly, we identified novel back-splicing junctions involving exon-intron boundaries or occurring entirely within intronic regions (Fig.1d, Supplementary Table S2). These events arise from the activation of cryptic splice sites within introns upon TDP-43 loss, which are subsequently utilized in back-splicing reactions to generate circRNAs containing CEs. circADGRL3 contains a CE that circularizes with a downstream annotated exon, generating an exon-intron circRNA composed of both cryptic and canonical sequences (Fig. 1g). Additional validated examples of such CE-canonical exon hybrid circRNAs include circPBX1, circFOXK1, circZNF423 (Supplementary Fig. 1d).

There are also circRNAs composed entirely of CEs arising from intronic regions derepressed upon loss of TDP-43. circGRM7 originates exclusively from three adjacent unannotated CEs within a single intron without incorporation of annotated exons (Fig.1h). Similarly, circDAPK1 consists of two CEs within intron 2 (Supplementary Fig. 1e). circUNC5Ds arise from a cluster of six unannotated CEs within intron 1, with distinct back-splicing junctions between different CEs generating multiple circUNC5D isoforms containing different combinations of these CEs (Supplementary Fig. 1e). Notably, these intronic cryptic circRNAs preferentially originated from significantly longer introns compared to other introns within the same host genes (Supplementary Fig. 1f), raising the possibility that extended intronic regions provide a permissive structural environment for aberrant back-splicing and cryptic circRNA formation upon TDP-43 loss. The full-length sequences of several circRNAs, in addition to their back-splicing junctions, were further validated by Sanger sequencing after cloning into plasmids (Supplementary Table S3).

Together, these findings reveal a previously unrecognized class of circRNAs regulated by TDP-43 and suggest that derepression of intronic splice sites contributes to aberrant circRNA biogenesis. We thereby refer to circRNAs that contain CEs and/or emerge de novo upon TDP-43 loss as cryptic circRNAs (Supplementary Table S2).

### A subset of cryptic circRNAs is associated with aberrant linear splicing upon loss of TDP-43 binding

To determine whether the upregulation of circRNAs is associated with altered TDP-43 binding, we profiled TDP-43–associated RNA regions in i^3^Neurons using Assay of Reverse Transcription-based RBP binding sites sequencing (ARTR-seq)^51^. ARTR-seq combines antibody-based targeting of RBPs with in situ reverse transcription, in which biotinylated dNTPs are incorporated into cDNA reverse-transcribed from RNAs proximal to the bound protein, thereby enabling enrichment and sequencing of RBP-associated RNA regions. Using an antibody against TDP-43, ARTR-seq identified widespread TDP-43–associated RNA regions in control neurons (Supplementary Fig. 2a-b). Analysis of enriched regions revealed significant enrichment of UG-rich motifs and predominant localization within intronic regions, consistent with known TDP-43 binding preferences (Fig. 2a, Supplementary Fig. 2b). In addition, TDP-43-associated transcripts identified by ARTR-seq showed substantial overlap with previously published TDP-43 eCLIP-seq datasets generated in K562 cells^52^, further supporting the specificity of the assay (Supplementary Fig. 2c). Notably, compared with circRNAs that were not significantly changed, upregulated circRNAs exhibited increased TDP-43 binding in intronic regions flanking their back-splice junctions (Fig. 2b). For example, ARTR-seq identified TDP-43–associated regions in the upstream introns of cryptic circPBX1 and circUNC13A, the downstream intron of circUNC5D.1, and both upstream and downstream flanking introns of circDIP2C (Fig. 2c).

**Figure 2.**
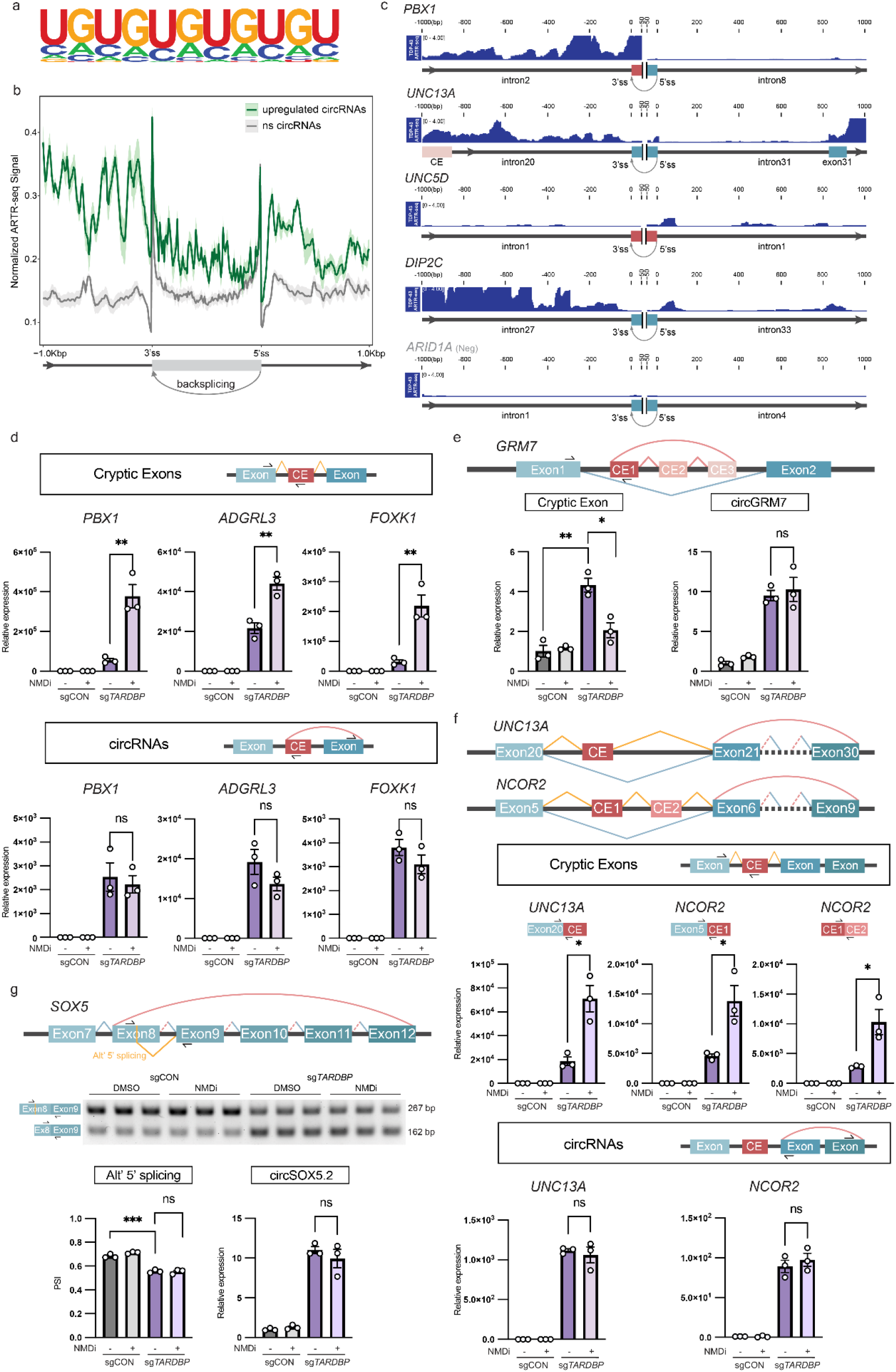
Upregulated circRNAs are associated with TDP-43 binding enrichment and are coupled to aberrant linear splicing. **a,** Motif analysis of TDP-43 ARTR-seq peaks reveals enrichment of the canonical UG-rich RNA-binding motif genome-wide. **b,** Distribution of TDP-43 binding across intronic regions flanking the back-splice junctions (±1 kb) of upregulated circRNAs (green) and non-significantly changed (ns) circRNAs (grey). Color shades indicate the standard error of the converge of three replicates. **c,** Representative TDP-43 binding profiles identified by ARTR-seq at the circPBX1, circUNC13A, circUNC5D.1 and circDIP2C loci. circARID1A (back-splice junction: chr1:26729651-26732792), an unchanged circRNA, is shown as a negative control. The horizontal axis represents ±1 kb of genomic sequence flanking the circRNA back-splice junction, centered in each panel. The 5′ and 3′ splice sites (ss) used in back-splicing are indicated. Exons are shown in light blue, introns in red, and CEs in pink. Dark-blue tracks represent normalized TDP-43 ARTR-seq signal. Black arrows indicate transcriptional orientation. **d–g,** Representative examples of coupling between cryptic circRNA biogenesis and aberrant splicing events induced by TDP-43 depletion following NMD inhibition. **d,** CE-containing linear transcripts and corresponding cryptic circRNAs. **e,** Intronic cryptic circRNA formation from adjacent CEs. **f,** Exonic circRNAs coupled to activation of neighboring cryptic splice sites. **g,** circRNA formation associated with alternative 5′ splice-site usage. Representative events were validated by RT–qPCR (d, e, f) or RT–PCR (g). Rounded arcs indicate back-splicing events, whereas angular arcs indicate linear splicing events. Blue arcs denote canonical splice junctions, orange arcs denote aberrant splice junctions induced by TDP-43 depletion, and red arcs denote cryptic circRNA-associated junctions. Bi-colored blue–red arcs indicate splice junctions shared between canonical and circular transcripts. Solid arcs represent experimentally validated or canonical splice junctions, whereas dashed arcs indicate predicted splice junctions. Arrows indicate primer positions used for RT–PCR or RT–qPCR analyses. Data are mean ± SEM from three biological replicates. Statistical significance was determined by two-tailed unpaired t-tests. ns, not significant. *P < 0.05, ** P < 0.01, *** P < 0.001, **** P < 0.0001.

Given the established role of TDP-43 in regulating alternative splicing of linear RNAs, we next examined the relationship between upregulated circRNA formation and aberrant splicing. To maximize detection of all aberrant transcripts, we incorporated datasets generated under both basal conditions and nonsense-mediated decay (NMD) inhibition following TDP-43 depletion^53^. Analysis of global mis-splicing events revealed that among the 569 non-intergenic upregulated circRNAs, 97 shared splice sites or were coupled with aberrant splicing events, including exon skipping, CE inclusion, and alternative splice site usage. To further characterize this relationship, we examined representative examples associated with distinct classes of splicing abnormalities.

We first focused on the CE-intronic circPBX1, circADGRL3 and circFOXK1 that incorporate both cryptic and canonical exons into their circularized sequences (Fig. 2d). RT-qPCR validation showed that loss of TDP-43 increased CE inclusion in the corresponding linear RNA, which was further markedly elevated following inhibition of NMD (Fig. 2d). This is consistent with previous observations that many CE–containing linear RNAs are preferentially degraded through NMD pathways^53, 54^. In contrast, the corresponding cryptic circRNAs were robustly detected upon TDP-43 knockdown and showed minimal changes following NMD inhibition (Fig. 2d), indicating that these circular isoforms evade NMD-mediated surveillance. These results suggest that cryptic splice sites activated upon TDP-43 loss can be used in both linear and circular RNAs. While linear transcripts containing CEs are continuously degraded by NMD, the corresponding circular products persistently accumulate.

We also examined intronic cryptic circRNAs that appeared to arise entirely from unannotated intronic sequences (Fig. 2e). As previously shown in Fig.1h, circGRM7 contained three adjacent CEs embedded within a single intron. Although corresponding linear CE products were not identified by the RNA-seq analysis, targeted RT-qPCR identified low-abundance linear CE transcripts upon TDP-43 depletion (Fig. 2e). Unexpectedly, these linear CE products were not stabilized by NMD inhibition and instead showed a modest reduction following SMG1 inhibition. This suggests that the low abundance of the linear isoform is likely due to preferential utilization of these cryptic splice sites for back-splicing and circRNA production, rather than active degradation of the linear CE-containing isoform.

Interestingly, some exonic circRNAs were associated with activation of neighboring cryptic splice sites without direct incorporation of the CE itself (Fig. 2f). For example, circUNC13A utilized a back-splice junction between exon 30 and 21, adjacent to a well-characterized CE inclusion event within intron 20 of UNC13A induced by TDP-43 loss^17, 18^. Thus, the 3’ splice site at the intron 20-exon 21 boundary is shared between the linear RNA and circRNA isoforms. Upregulation of the CE-containing linear RNA was accompanied by increased expression of circUNC13A (Fig. 2f). This suggests that the binding of TDP-43 in intron 20 (Fig. 2c) suppresses both cryptic splice sites usage and back-splicing, resulting in coordinated regulation of the linear and circular isoforms. Consistent with previous observations, the CE-containing linear transcript was actively degraded and showed a marked increase following NMD inhibition, whereas the circUNC13A was not affected (Fig. 2f). Likewise, circNCOR2 contains a back-splice junction between exons 9 and 6, and its upregulation was coupled with activation of two CEs within intron 5 in the corresponding linear transcript, which is also subject to NMD-mediated degradation (Fig. 2f). Together, these findings suggest that altered splice site selection following TDP-43 depletion can broadly alter back-splicing patterns beyond direct CE circularization.

We next investigated whether other forms of aberrant splicing were also coupled with circRNA dysregulation. In the SOX5 transcript, TDP-43 depletion induced a switch toward usage of an alternative 5′ splice site within exon 8, located adjacent to the circSOX5.2 back-splice junction, resulting in an isoform with a shortened exon 8 (Fig. 2g). RT-PCR confirmed decreased full-length exon 8 and increased shortened exon 8 upon TDP-43 knockdown. Unlike CE– containing linear transcripts, this alternative splice isoform was not substantially altered by NMD inhibition (Fig. 2g). circSOX5.2 was similarly increased following TDP-43 depletion and remained largely unaffected by NMD inhibition, further supporting a mechanistic coupling between aberrant splice site selection and circRNA biogenesis that extends beyond canonical NMD-sensitive CEs.

Together, these findings demonstrate a strong coupling between aberrant linear splicing and cryptic back-splicing following TDP-43 loss of function in a subset of transcripts. These results suggest that TDP-43 binding can influence multiple aspects of splice site selection, leading to coordinated regulation of distinct RNA isoforms, including both linear and circular RNA species. Activation of derepressed splice sites can promote the generation of both linear and circular aberrant RNA isoforms, yet the resulting products exhibit markedly different fates. Whereas many cryptic linear transcripts are selectively degraded by RNA quality control pathways such as NMD, cryptic circRNAs evade decay, highlighting their potential as durable molecular readouts of TDP-43 dysfunction.

### Cryptic circRNAs display increased stability and accumulate progressively in neurons

Beyond the differential effects of NMD on cryptic linear transcripts, circRNAs are generally more stable than linear RNAs due to their resistance to exonucleolytic degradation. To further characterize the properties of cryptic circRNAs, we examined their stability relative to the corresponding canonical linear transcripts in TDP-43 knockdown neurons. RNA half-life was measured following transcriptional shutoff induced by actinomycin D treatment. While the linear mRNA gradually decreased, circRNAs exhibited minimal decay over the measured time course, indicating greater stability than their linear counterparts (Fig. 3a).

**Figure 3.**
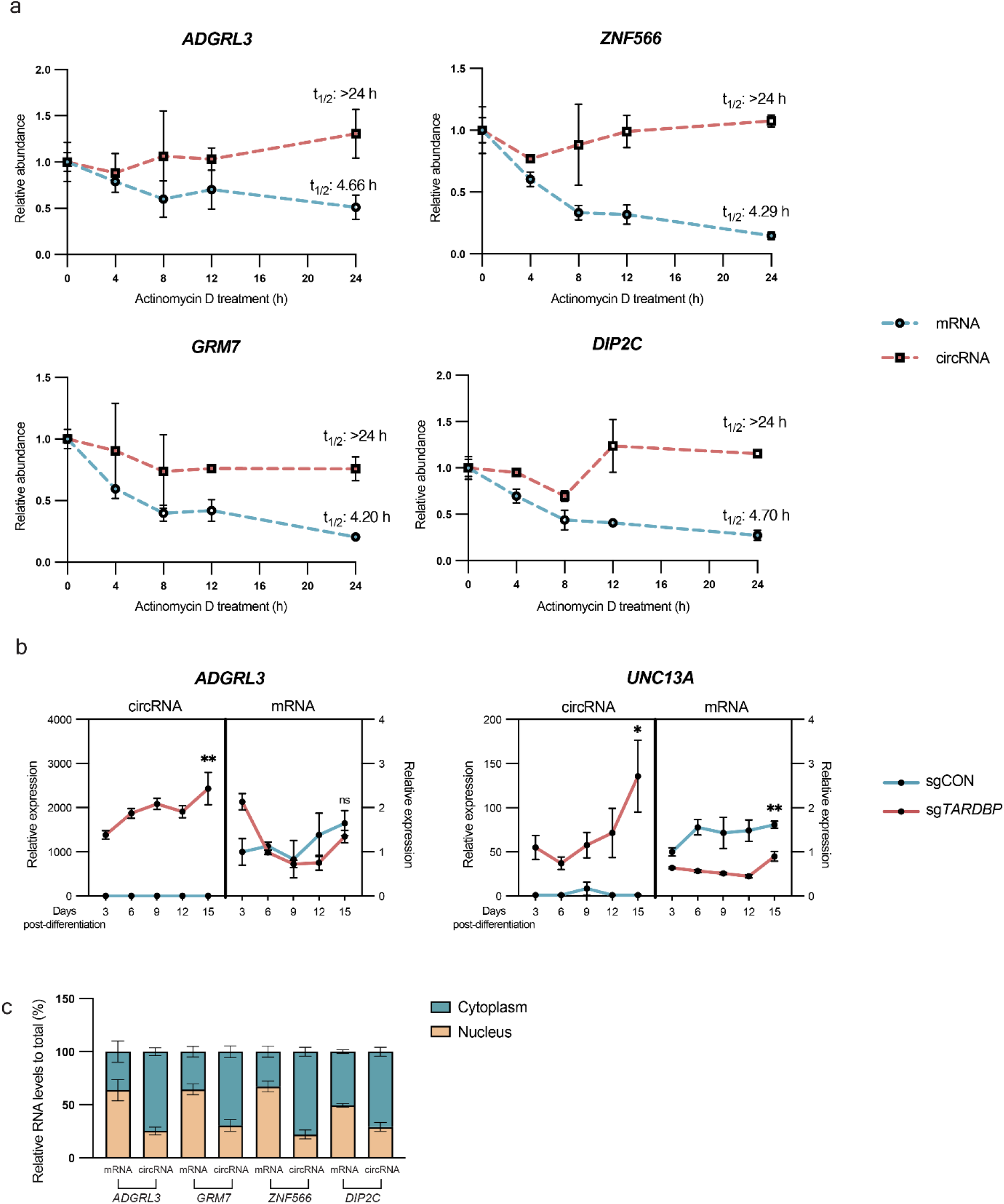
Cryptic circRNAs are stable and accumulate progressively in neurons. **a,** RNA decay analysis of representative circRNAs and corresponding host mRNAs in TDP-43 knockdown i^3^Neurons. RNA levels were measured by RT–qPCR at indicated time points following actinomycin D treatment and normalized to the RNA spike-in control. All time points were compared to time 0. Estimated transcript half-lives (t₁/₂) are indicated. Data are mean ± SEM from two biological replicates. **b,** RT–qPCR analysis of circRNAs and corresponding host mRNAs during neuronal differentiation in control and TDP-43 knockdown i^3^Neurons. RNA levels were measured at the indicated days post-differentiation and normalized to HPRT1, which remains stable post-differentiation^50^. Data are mean ± SEM from three biological replicates. Statistical significance was determined for day 15 samples using two-tailed unpaired t-tests. ns, not significant; *P < 0.05, ** P < 0.01. **c,** RT–qPCR analysis of circRNAs and corresponding host mRNAs in cytoplasmic and nuclear fractions from TDP-43 knockdown i^3^Neurons. RNA levels in each fraction are normalized to the RNA spike-in control and presented relative to total RNA. Data are mean ± SEM from four biological replicates.

We next asked whether this difference in stability is reflected in RNA dynamics during neuronal differentiation. Time-course analysis showed that cryptic circADGRL3 and circUNC13A increased over time in differentiated neurons with constitutive TDP-43 knockdown, while their mRNAs remained largely unchanged (Fig. 3b, Supplementary Fig. 3a). These observations suggest that linear transcripts remain at a relatively stable level in neurons, whereas their circular isoforms persist and progressively accumulate over time.

Next, we examined the subcellular distribution of circRNAs and their corresponding linear transcripts. Biochemical fractionation was performed to separate nucleus and cytoplasm from TDP-43-depleted neurons, which was validated by known nuclear- and cytosolic-enriched transcripts (Supplementary Fig. 3b). RT-qPCR analysis of selected cryptic circRNAs and their corresponding mRNAs revealed that circRNAs were predominantly localized to the cytoplasm and, in many cases, exhibited greater cytoplasmic enrichment than their host mRNAs (Fig. 3c).

### Cryptic circRNAs are elevated in human postmortem brain tissues with TDP-43 proteinopathy

To determine whether TDP-43–regulated circRNAs are altered in human disease tissues, we measured candidate circRNAs in postmortem temporal cortex from individuals with ALS/FTD and non-neurological controls. We examined both C9ORF72-ALS/FTD and sporadic FTD cases. RT–qPCR analysis of candidate cryptic circRNAs revealed significant upregulation, with circADGRL3 and circUNC13A showing the most robust increases across both C9ORF72 ALS/FTD and sporadic FTD cases compared with controls (Fig. 4a). In contrast, the corresponding host mRNAs exhibited only modest or no significant changes (Supplementary Fig. 4a), suggesting that the circRNA increase is not due to altered transcription of the host genes. Notably, the magnitude of circRNA induction was comparable to, and in some cases greater than, established linear CE–containing transcripts, including UNC13A, CAMK2B, STMN2 and HDGFL2 (Fig. 4b). As TDP-43 proteinopathy is also found in Alzheimer’s disease (AD), we examined the expression of those target circRNAs along with their host mRNAs in the same brain region of AD patients’ postmortem tissues. CircRNAs exhibited substantially more pronounced changes than their linear host transcripts in AD tissues (Fig. 4c, Supplementary Fig. 4b), and displayed expression patterns distinct from those observed in ALS/FTD (Fig. 4a, c). While circADGRL3 and circUNC13A showed robust upregulation in ALS/FTD (Fig. 4a), they were largely undetectable in AD samples (Fig. 4c). In contrast, circZNF566, circDIP2C and circNCOR2 exhibited more pronounced upregulation in AD samples, while showing weaker or no significant changes in ALS and FTD (Fig. 4a, c). Linear CEs also displayed distinct disease-associated patterns. For example, UNC13A CE showed the highest upregulation in ALS/FTD while it was not detectable in AD, whereas CAMK2B CE exhibited greater elevation in AD than in ALS/FTD (Fig. 4b, d). These results suggest that splicing dysregulation downstream of TDP-43 dysfunction likely exhibits disease-specific patterns across neurodegenerative diseases with TDP-43 proteinopathy, although more cases need to be analyzed to confirm these observations.

**Figure 4.**
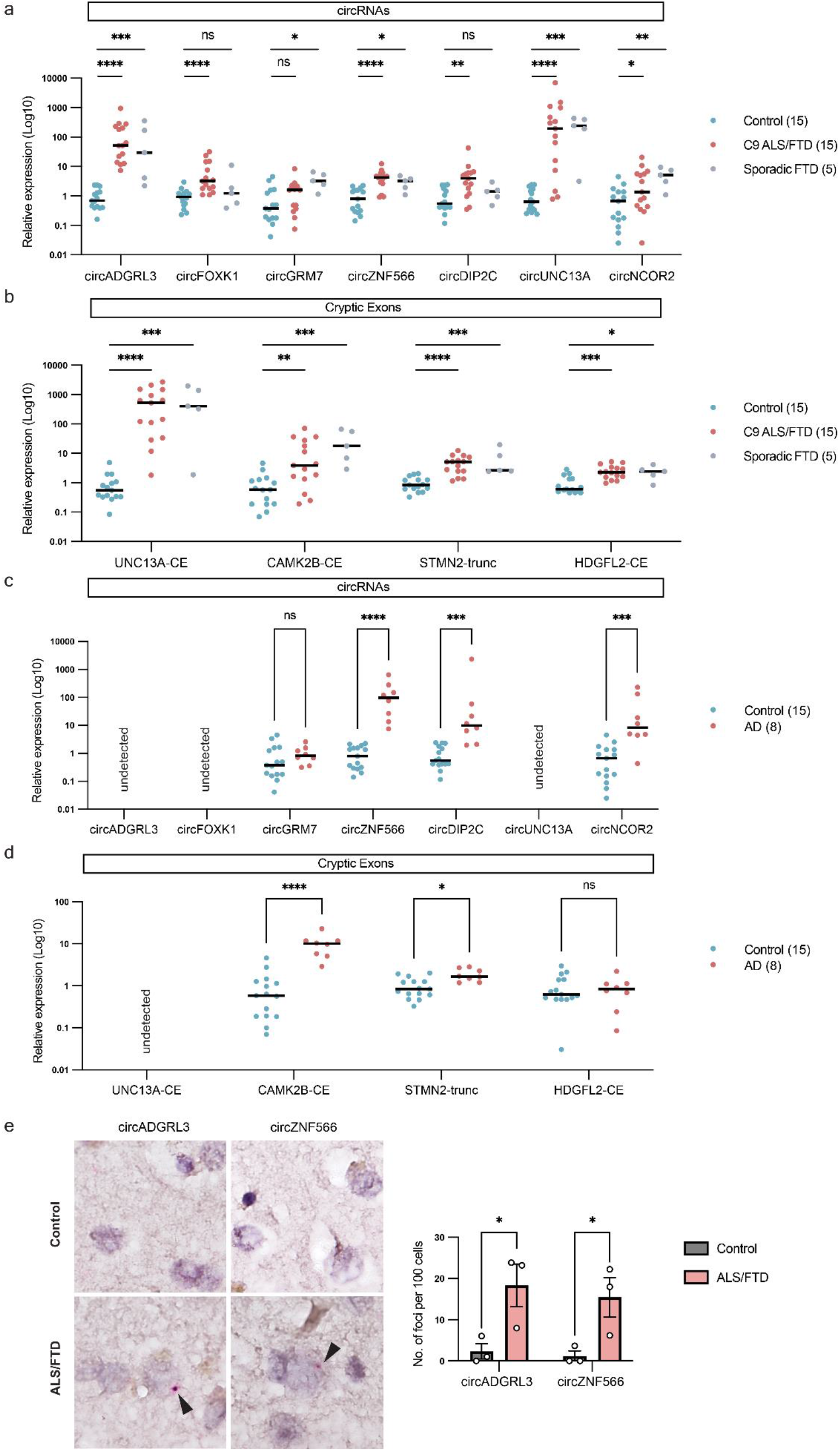
Cryptic circRNAs are elevated in postmortem brain tissues from patients. **a–d,** RT–qPCR analysis of cryptic circRNAs (a, c) and linear CE–containing transcripts (b, d) in temporal cortex tissues from non-neurological controls (n = 15), C9ORF72-associated ALS/FTD (n = 15), sporadic FTD cases (n = 5) (a, b), and Alzheimer’s disease (AD; n = 8) (c, d). Relative expression levels were calculated normalized to HPRT1 and are shown on a log10 scale. Each dot represents a sample from an individual donor, and horizontal bars indicate median values. Statistical significance was determined using two-tailed Mann–Whitney tests. ns, not significant. *P < 0.05, ** P < 0.01, *** P < 0.001, **** P < 0.0001. **a,** Expression of seven candidate cryptic circRNAs in control, C9ORF72-associated ALS/FTD, and sporadic FTD temporal cortex tissues. **b,** Expression of representative linear CE–containing transcripts, including UNC13A CE, CAMK2B CE, truncated STMN2 (STMN2-trunc), and HDGFL2 CE, measured in the same ALS/FTD cohort. **c,** Expression of seven candidate cryptic circRNAs in control and AD temporal cortex tissues. “Undetected” indicates transcripts that were not reliably detected in the analyzed samples. **d,** Expression of representative linear CE–containing transcripts in control and AD temporal cortex tissues. “Undetected” indicates transcripts that were not reliably detected by RT-qPCR in the analyzed samples. **e,** BaseScope in situ hybridization targeting the back-splice junctions of circADGRL3 and circZNF566 in temporal cortex tissues from control and ALS/FTD cases. Representative images are shown. Arrowheads indicate positive BaseScope signals. Quantification of circRNA-positive puncta per 100 cells is shown on the right. Each dot represents an individual donor (n = 3 per group), and bars indicate mean ± SEM. Statistical significance was determined using two-tailed unpaired t-tests. *P < 0.05.

We further validated the upregulation of cryptic circRNAs in human brain tissue using BaseScope in situ hybridization with probes targeting the back-splice junctions of circADGRL3 and circZNF566. In control samples, signals were barely detectable. In contrast, ALS/FTD brains exhibited clear BaseScope signals (Fig. 4e, arrowheads). Quantification revealed a significant increase in the number of puncta for both circRNA targets in disease samples compared to controls (Fig. 4e), further supporting the robust upregulation of cryptic circRNAs in ALS/FTD.

Together, these results demonstrate that cryptic circRNAs are elevated in postmortem human brain tissues with TDP-43 proteinopathy and exhibit disease-associated, and in some cases disease-selective, expression patterns.

## DISCUSSION

In this study, we identified a novel role for TDP-43 in regulating back-splicing and showed that loss of TDP-43 binding is associated with the emergence of a distinct class of circular RNAs termed cryptic circRNAs. While TDP-43 has been well established as a regulator of canonical linear splicing, our findings extend this framework by demonstrating that TDP-43 also constrains circular RNA biogenesis (Fig. 5), thereby linking TDP-43 proteinopathy to the generation of stable, noncoding RNA species.

**Figure 5.**
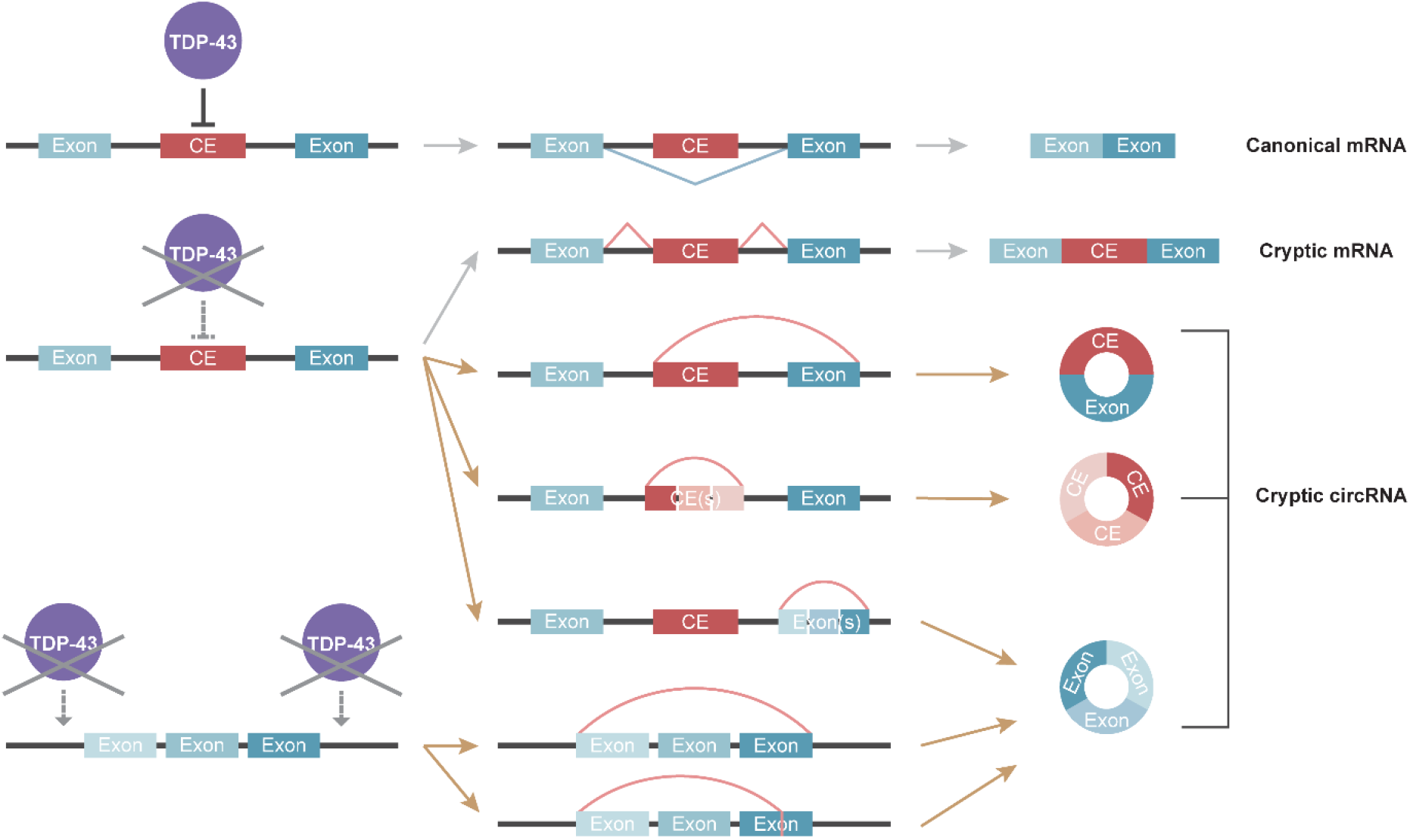
Loss of TDP-43 unleashes cryptic circRNA biogenesis. Under normal conditions (top), TDP-43 binding suppresses CE inclusion and back-splicing, allowing canonical mRNA production. Upon loss of TDP-43 (bottom), derepressed cryptic splice sites are utilized to generate linear CE-containing mRNA isoforms and/or exploited in back-splicing to produce cryptic circRNAs, including exon-intronic circRNAs (incorporating both CE and canonical exon sequences), intronic circRNAs (composed entirely of CE sequences), and exonic circRNAs (composed of canonical exons). Back-splicing of exonic circRNAs could be coupled to activation of neighboring cryptic splice sites, or independent of CEs.

Under normal conditions, TDP-43 binds near cryptic splice sites and represses CE inclusion. Upon loss of TDP-43, these regions become permissive for aberrant splice-site selection, leading to incorporation of CEs into linear mRNAs. A subset of these derepressed splice sites can also undergo back-splicing, resulting in incorporation of CEs into circular RNAs. Some cryptic circRNAs contain both cryptic and canonical exons, whereas others are composed entirely of intronic-derived cryptic sequences. Beyond these CE-containing species, loss of TDP-43 also enhances back-splicing of canonical exons, with the majority of upregulated circRNAs comprising only canonical exons. Some of these circRNAs arise independently of detectable changes in the corresponding linear transcripts, whereas others are associated with nearby alternative splicing events. While some are detectable at baseline and further increased upon TDP-43 depletion, others arise de novo following TDP-43 loss. Together, these observations indicate that TDP-43 suppresses circular RNA biogenesis at both cryptic and canonical exons. The determinants governing circRNA enhancement and the preference toward circular versus linear cryptic splicing remain unclear. Features such as splice-site strength, intronic architecture, RNA secondary structure, transcriptional kinetics, and additional RNA-binding proteins may contribute. Defining how these factors shape transcript-specific responses will be important for understanding RNA dysregulation downstream of TDP-43 loss.

Beyond their biogenesis, cryptic circRNAs possess molecular properties that favor their persistence. For CE-containing RNA isoforms, the linear transcripts are often degraded by NMD and related RNA quality control pathways, limiting their accumulation, whereas cryptic circRNAs are generally not subjected to NMD-mediated decay. Combined with the intrinsic resistance of circRNAs to exonucleolytic degradation, these properties allow cryptic circRNAs to persist and accumulate as durable molecular readouts of splicing dysregulation. Notably, circRNAs have also been shown to be preferentially packaged into extracellular vesicles (EVs), such as exosomes^55, 56^. These features highlight the potential of cryptic circRNAs as accessible biomarkers of TDP-43 dysfunction.

The detection of cryptic circRNAs in postmortem human brain tissues supports their relevance to neurodegenerative diseases associated with TDP-43 proteinopathy. We observed disease-specific expression patterns, with circADGRL3 and circUNC13A preferentially increased in ALS and FTD, whereas circZNF566, circDIP2C, and circNCOR2 showed more pronounced elevation in AD. These differences may reflect heterogeneity in the distribution, timing, or co-factors of TDP-43 pathology across disorders and suggest that distinct cryptic circRNAs may capture different manifestations of TDP-43 dysfunction. However, larger cohorts are required to systematically define circRNA signatures across disease stages, subtypes, and brain regions. In addition, analysis of biofluid samples will be essential to determine whether selected circRNAs can serve as accessible biomarkers of TDP-43 loss of function for disease diagnostic and prognostic applications.

Together, these findings expand the scope of TDP-43–dependent RNA regulation to back-splicing and identify cryptic circRNAs as stable molecular consequences of TDP-43 loss of function in human neurodegeneration. By uncovering a pathway connecting cryptic splice site activation to circRNA biogenesis, this work provides a new framework for understanding RNA dysregulation and its translational potential.

## ACKNOWLEDGMENTS

We thank Johns Hopkins Brain Resource Center for providing postmortem human brain tissue samples. We thank all the Sun lab members for the support. This work is supported by R01NS127925 (S.S.), R01AG078948 (S.S.), RF1NS113820 (S.S.), and RM1 008935 (C.H.) from NIH, and Robert Packard Center for ALS Research. Z.Z. was a recipient of the Milton Safenowitz Post-Doctoral Fellowship from the ALS Association, the Toffler Scholar Award and the Postdoc Development Grant from Muscular Dystrophy Association (MDA). Y.Y. received the Toffler Scholar Award. N.X. is a recipient of Maryland Stem Cell Research Post-Doctoral Fellowship. C.H. is an investigator of the Howard Hughes Medical Institute.

## AUTHOR CONTRIBUTIONS

C.Z. and S.S. conceived the study, designed the experiments, and wrote the manuscript. C.Z. performed the majority of molecular and cellular biology experiments and led experimental design and data interpretation. Z.Z. (Zhiyan Zhao) conducted bioinformatic analyses and contributed to selected molecular experiments. B.S. assisted with cell culture and molecular experiments. Y.C. and Y.X. contributed to ARTR-seq library construction and analysis under the mentorship of C.H. Z.Z. (Zhe Zhang) also contributed to ARTR-seq library preparation and assisted with cell culture. Y.Y. generated the sgRNA iPSC cell lines. N.X. and R.Z. performed BaseScope experiments and analysis. Y.H. processed human patient tissue samples. J.C.T. provided human postmortem tissue samples from the BRC. S.S. supervised the study.

## METHODS

### Plasmids

Single guide RNAs (sgRNAs) were cloned into the pU6-sgRNA EF1α-puro-T2A-BFP vector (Addgene plasmid #60955)^50^. The sgRNA sequence used were as follows: non-targeting control: GGACTAAGCGCAAGCACCTA, and TDP-43: GGCCCGCGCGTGCCAGCCGA.

For sequence validation of circRNAs, divergent primers (Supplementary Table S4) were used to amplify circRNA-derived cDNA using 2 × Phanta UniFi Master Mix (Dye Plus) (Vazyme, #P526) with 32 PCR cycles. PCR products were resolved on 2.5% agarose gels, and bands of the larger size that contain the complete circRNA sequences were excised and purified using a Gel Extraction Kit (Thermo Fisher Scientific, #K0692). Purified amplicons were cloned into plasmids using the Zero Blunt TOPO PCR Cloning Kit (Invitrogen, #450245) according to the manufacturer’s instructions, and positive clones were subjected to Sanger sequencing using universal vector primers.

### Cell culture and transfection

HEK293T cells were maintained in Dulbecco’s Modified Eagle Medium (DMEM) supplemented with 10% (v/v) fetal bovine serum (FBS), 100 U/mL penicillin, and 100 μg/mL streptomycin at 37°C in a humidified incubator with 5% CO₂. HEK293T cells were used for lentiviral packaging of sgRNA constructs following standard protocols as previously described^31^.

### Human iPSC and iPSC-derived neurons

Human iPSCs and their differentiation into i^3^Neurons were performed as previously described^57^. Briefly, engineered iPSCs expressing dCas9 and doxycycline-inducible mNGN2 were maintained on Matrigel-coated plates (Corning, #354277) in Essential 8 medium (Thermo Fisher Scientific, #A1517001) and passaged with Accutase (Innovative Cell Technologies, #AT104-500) at ∼80% confluency. Neuronal differentiation was initiated by dissociation and culture in N2 induction medium (ThermoFisher Scientific, #12660012, #17502048, #1140050) supplemented with doxycycline (Sigma-Aldrich, #D9891) for 3 days, followed by replating onto poly-L-ornithine–coated (Sigma-Aldrich, #P3655) plates in BrainPhys-based neuronal differentiation medium (StemCell Technologies, # 05790) containing B27 (Thermo Fisher Scientific, #17504044), NT-3 (Thermo Fisher Scientific, #450-03), BDNF (Thermo Fisher Scientific, #450-02), laminin (Thermo Fisher Scientific, #L2020), and doxycycline. Half-medium changes were performed every other day, and neurons were harvested at day 14 unless otherwise indicated.

For ARTR-seq experiments, the PiggyBac-TO-hNGN2 iPSC line was generated as previously described^58^. Briefly, iPSCs were obtained from The Jackson Laboratory (#JIPSC001104) and co-transfected with PB-TO-hNGN2 and pEF1α::Transposase (gifts from Dr. Michael Ward) using Lipofectamine Stem Transfection Reagent (Thermo Fisher Scientific, #STEM00001), followed by puromycin selection. Neuronal differentiation was performed using conditions similar to those described for i^3^Neurons, with minor modifications. Cells were maintained in N2 induction medium for an additional day, during which proliferating cells were eliminated by treatment with 5-fluoro-2′-deoxyuridine (MedChemExpress, #HY-B0097) and uridine (StemCell Technologies, #100-0541). Neurons were then cultured in the same BrainPhys-based differentiation medium used for i^3^Neurons with the addition of GDNF (1 μg/mL, Thermo Fisher Scientific, #450-10). Half of the medium was replaced every other day, and cells were harvested at day 14.

### i^3^Neurons treatments and processing

For NMD inhibition experiments, i^3^Neurons were treated with 0.5 μM hSMG1 inhibitor (MCE, #HY-124719) for 24 hours prior to harvest, with DMSO used as a negative control.

For cytoplasmic and nuclear fractionation, i^3^Neurons were resuspended in NP-40 lysis buffer (10 mM Tris-HCl, pH 7.5; 10 mM NaCl; 0.2% NP-40; 1 U/μL RNase inhibitor) and incubated on ice for 5 minutes. One-third of the total lysate was collected as the input (total) fraction. Lysates were then centrifuged at 3,000 rpm for 7 minutes at 4 °C. The supernatant, corresponding to the cytoplasmic fraction, was transferred for RNA extraction. The nuclear pellet was washed three times with mild wash buffer (10 mM Tris-HCl, pH 8.0; 10 mM NaCl; 1 mM EDTA), with centrifugation at 3,000–5,000 rpm for 5 minutes at 4 °C between washes. After removal of the supernatant, the nuclei were collected for RNA extraction.

### Human brain tissue source and processing

Frozen and formalin-fixed paraffin-embedded (FFPE) human postmortem brain tissues were obtained from Johns Hopkins Brain Resource Center. Subject demographics are listed in Table S5. Usage of human samples and data was approved by the Johns Hopkins University School of Medicine Office of Human Subjects Research Institutional Review Boards.

Frozen human brain tissue was sectioned into 50 mg pieces and homogenized using an Omni Tissue Master 125 Homogenizer in 250 μL of ice-cold homogenization buffer containing 1× TE, 2 mM PMSF, 1× protease inhibitor cocktail (Sigma-Aldrich, #11697498001), and 1× phosphatase inhibitor cocktail (Bimake, #B15002). Aliquots of homogenate were processed and immediately used for RNA extraction.

### RNA extraction, RT-PCR and RT-qPCR

Total RNA was extracted from cultured cells and human postmortem tissues using TRIzol Reagent (Thermo Fisher Scientific, #15596026) according to the manufacturer’s instructions. Complementary DNA (cDNA) was synthesized using HiScript III RT SuperMix for qPCR (+gDNA wiper) (Vazyme, #R323). For RNA half-life, subcellular fractionation and RNA accumulation experiments, equal amounts of C*Luc* RNA spike-in control (NEB, #E1610S) were added to each normalized RNA sample prior to reverse transcription and subsequently used as an external reference for normalization.

Semi-quantitative PCR was performed using 2× Rapid Taq Master Mix (Vazyme, #P222-03) with 30–33 amplification cycles to assess splicing changes. PCR products were resolved on 2.5% agarose gels and imaged using a ChemiDoc Imaging System (Bio-Rad). Relative band intensities were quantified with background subtraction using the Image Lab software (Bio-Rad). Percent spliced in (PSI) values were calculated based on the relative intensities of the long (full-length) and short isoform PCR products using the formula: PSI = long / (long + short).

Quantitative PCR (qPCR) was performed in technical duplicates for each cDNA sample using Taq Pro Universal SYBR qPCR Master Mix (Vazyme, #Q712) on QuantStudio 6 and QuantStudio 3 Flex Real-Time PCR Systems (Thermo Fisher Scientific). Relative expression levels were calculated using the ΔΔCt method. *CLuc* spike-in RNA was used for normalization in RNA half-life, subcellular fractionation, and RNA accumulation experiments; GAPDH was used as the reference gene for NMD inhibition experiments; and HPRT1 was used as the reference gene for all other qPCR analyses. For targets with undetermined Ct values, a Ct value of 40 was assigned for downstream statistical analysis and visualization, corresponding to the maximum number of amplification cycles. Primer sequences are provided in Table S4 and references for previously published primers are indicated in the Source column of the table^18, 59–62^.

### RNA half-life measurement

i^3^Neurons were treated with 3 μg/mL actinomycin D (Sigma-Aldrich, #A9415) and harvested at 0, 4, 8, 12, and 24 h prior to harvest at day 14. Total RNA was isolated at each time point and analyzed by RT–qPCR as described above. RNA relative abundance was normalized to the 0 h time point for each transcript. Transcript half-lives were estimated by nonlinear regression using a one-phase exponential decay model in GraphPad Prism software. Half-lives were successfully calculated for linear mRNAs. For circRNAs, transcript abundance remained largely unchanged throughout the 24 h time course and could not be reliably fitted by the decay model; therefore, their half-lives were considered to exceed the experimental observation period (>24 h).

### circRNA-related RNA-seq library preparation

For each sample, 5 μg of total RNA isolated from i^3^Neurons was used for circRNA-enriched library preparation. Poly(A)+ mRNA was first isolated using the NEBNext Poly(A) mRNA Magnetic Isolation Module (NEB, #E7490L) according to the manufacturer’s instructions. The poly(A)(–) fraction was collected as the flowthrough from the oligo(dT)\₍₂₅₎ bead selection step and subsequently purified using the Quick-RNA Microprep Kit (Zymo, #R1050). The purified poly(A)(–) RNA was then treated with 5 U RNase R (NEB, #M0100S) at 37 °C for 20 min to enrich for circular RNAs. Following digestion, RNA was recovered using TRIzol reagent and quantified using a Qubit fluorometer. For downstream processing, 500 ng of poly(A)(–), RNase R–treated RNA was subjected to rRNA depletion using the Ribo-off rRNA Depletion Kit (Vazyme, #N406). The resulting poly(A)(–)/RNase R(+)/rRNA(–) circRNA-enriched RNA was immediately used for RNA-seq library preparation with the VAHTS Universal V10 RNA-seq Library Prep Kit for Illumina (Vazyme, #NR606).

In parallel, 50 ng of the corresponding poly(A)+ mRNA fraction was processed for library construction using the same kit (Vazyme, #NR606). Three independent biological replicates were prepared for each group. Libraries were sequenced on the Illumina NovaSeq X Plus platform to generate 150-bp paired-end reads.

### RNA-seq analysis

#### Genome reference

The *Homo sapiens* reference genome (GRCh38.p14; Ensembl release 112) was used for all analyses, together with GENCODE v32 gene annotation. Reference sequences for ribosomal RNAs (rRNAs), including 28S (NR_003287.4), 18S (NR_003286.4), 5.8S (NR_003285.3), and 5S rRNA, were downloaded from the National Center for Biotechnology Information (NCBI) Reference Sequence database.

#### Gene expression analysis

Raw reads of the poly(A) library were trimmed using Cutadapt version 5.0^63^ and filtered for quality (Phred Score >= 33). Processed reads were mapped using HISAT2 version 2.2.1^64^, deduplicated using Picard MarkDuplicates version 3.3.0 (Broad Institute), then quantified using FeatureCounts version 2.0.8^65^. Highly expressed genes (>=10 reads in all samples) were used for differential expression analysis. Filtered read counts were normalized using DESeq2 (version 1.46.0)^66^ with size factor normalization, and differential expression analysis was subsequently conducted. Log2FoldChange and p-adjusted values were used to determine differentially expressed genes (|LFC|>=1, p-adjusted < 0.05). Mapped reads were visualized using deepTools bamCoverage version 3.5.6^67^.

#### Detection and analysis of circRNAs

Raw reads of the poly(A)(–)/RNase R(+)/rRNA(–) library were trimmed using Cutadapt version 5.0^63^ and filtered for quality (Phred Score >=33). Back-splicing junctions (BSJs) were detected using CIRI2^68^ using the default parameters described by the official documentation. High confidence circRNAs (BSJs >=5 in at least 50% of samples) were used for differential expression analysis. Filtered BSJ counts were normalized using DESeq2 (version 1.46.0)^66^ with size factor normalization, and differential expression analysis was subsequently conducted. Log2FoldChange and p-values were used to determine differentially expressed genes (|LFC|>=0.5, p < 0.05). Linear cryptic junctions within circRNAs were detected using LeafCutter version 0.2.9^69^ from bam files.

#### Alternative splicing analysis

RNA-seq datasets generated from sgCON and sg*TARDBP* i^3^Neurons treated with or without nonsense-mediated decay (NMD) inhibition were obtained from Sinha, I.R.*, et al* (PRJNA1235234)^53^. Alternative splicing events were identified using rMATS (v4.3.0)^70^ with parameters --novelSS --variable-read-length --individual-counts -t paired. Events were retained with junction coverage ≥10, |ΔPSI| ≥ 0.1, and FDR < 0.05. Events were integrated across comparisons, and summary PSI values were defined as the maximum PSI across non-TDP conditions (max_PSI_control) and TDP plus NMD-factor perturbations (max_PSI_TDP_NMD). Differential splicing was calculated as dPSI_TDPvsCTRL, and events were classified as increased or decreased inclusion (|ΔPSI| > 0.1). Events absent in sgTDP were classified as “NMD”. Events were converted to BED format and intersected with circRNA back-splice junctions using BEDTools^71^ (bedtools intersect). Overlaps were refined by requiring exact matches between back-splice coordinates and annotated splice-site boundaries. Unique circRNA–event pairs were retained and summarized.

### ARTR-seq library preparation

ARTR-seq was performed as previously described^27, 31^, with minor modifications. Briefly, cells cultured in imaging chambers were fixed with 1.5% paraformaldehyde for 10 min, quenched with 125 mM glycine for 1 min, permeabilized with 0.5% Triton X-100 on ice for 10 min, and blocked in 0.1% BSA in DPBS supplemented with RNase inhibitor (Vazyme, #R301-03). TDP-43 was targeted using a TDP-43 antibody (Abnova, clone 2E2-D3, #H00023435-M01; 1:500), followed by Alexa Fluor 546 secondary antibody (ThermoFisher Scientific, # A11003) and pAG-RTase incubation. One sample was additionally stained with Hoechst (Cell Signaling Technology, #4082S) for nuclear counterstaining and imaged by confocal microscopy for validation. Input samples were processed in parallel without primary antibody. Reverse transcription was performed in situ using an adapter-RT primer (5′-AGACGTGTGCTCTTCCGATCTNNNNNNNNNN-3′) and a nucleotide mix containing biotin-16-dUTP and biotin-16-dCTP (Jena Bioscience, # NU-803-BIO16-S, #NU-809-BIO16-S). After terminating the reaction with 20 mM EDTA and 10 mM EGTA, samples were digested with proteinase K (Thermo Fisher Scientific, #AM2548), recovered by phenol-chloroform extraction and ethanol precipitation, treated with RNase H and RNase A/T1, and biotinylated cDNA was enriched using pre-blocked streptavidin beads (Thermo Fisher Scientific, #65001). A 3′ cDNA adapter (5′Phos-8N-AGATCGGAAGAGGTCGTGT-3′SpC3) was then ligated, and libraries were PCR amplified and gel purified to select fragments of 180– 400 bp for sequencing. Sequencing was performed with three biological replicates per group at the University of Chicago Genomics Facility on an Illumina NovaSeq 6000 platform with a paired-end read length of 150 bp.

### ARTR-seq analysis

The adapter sequences in the read were trimmed with Cutadapt version 5.0^63^ using the parameters: --nextseq-trim=20 --action=trim -m 20 -e 0.15 -n 3 -O 5 -a AGATCGGAAGAGCACACGTCT -A GATCGTCGGACTGTAGAACTCTGAAC; the 8 nt unique molecular identifier (UMI) sequences at the 5′ end of the Read1 (R1) and the 3′ end of the Read2 (R2) were removed. The UMI trimmed from R1 was added to the read name for further deduplication. An extra 4 nts at the 3′ end and the 5′ end of the R1 and R2, respectively were removed from the adapter-free sequence to minimize mapping mismatch caused by the imperfect paired sequence in the random primer.

The reads were first mapped to the corresponding rRNA sequences using Bowtie2 (v.2.4.4)^72^ with parameters: –seedlen=15. The mapped reads were discarded to remove rRNA contamination. The remaining unmapped reads were mapped to the corresponding genome using STAR (v2.7.11b) with parameters: --alignEndsType EndToEnd --outFilterMatchNmin 20 --alignMatesGapMax 15000 --outFilterMismatchNmax 10 --outFilterMultimapNmax 50 --outSAMmultNmax -1 --quantMode TranscriptomeSAM --outSAMtype BAM SortedByCoordinate --outFilterType BySJout --outSAMattributes NH HI AS nM NM MD jM jI MC --outReadsUnmapped Fastx. Uniquely mapped reads were deduplicated to get the usable reads using UMICollapse^73^ with the parameter: -t 2 -T 16 --paired --data naive --merge avgqual --two-pass. For visualization, .bam files of the usable reads were converted to bigWig with bamCoverage in the deepTools suite (v.3.5.5)^67^ with normalization by its respective sequencing depth using the parameters –normalizeUsing BPM–binSize 1. The ARTR-seq read profiles were calculated by computeMatrix from the deepTools suite and generated by in-house Rscript with ggplot2.

For peak calling, the usable reads in one library were first split into two .bam files containing reads aligned to the positive and negative strands, respectively. The paired-end .bam was converted to .bed for peak calling using bamtobed in the BEDTools suite with the parameters: -split. MACS3^74^ was used to identify peaks with default parameters, except for adding ‘–keep-dup -p BEDPE’. MACS3 gives the fold enrichment (signal value) and P value based on Poisson distribution, and corrects the P values for multiple comparison using the Benjamini–Hochberg correction. The peaks located in two strands were called separately using the corresponding strand read in the corresponding input libraries as background. The peak files for one library were later combined.

### Protein extraction and Immunoblotting

i^3^Neurons were detached by DPBS washes and collected by brief centrifugation. The cell pellets were lysed in RIPA buffer supplemented with 1× protease inhibitor cocktail (Sigma-Aldrich, #11697498001) and incubated on ice for 20 min, followed by centrifugation at 12,000 × g for 20 min at 4°C. The supernatant was collected, and protein concentration was quantified using BCA Assay (Thermo Fisher Scientific, #23225).

Equal amounts of protein (5 μg) were mixed with 5× loading buffer, heated at 90°C for 2 min, and separated on a 10% SDS–PAGE gel alongside a protein ladder (Bio-Rad, #1610374). Proteins were then transferred onto a nitrocellulose membrane using a wet transfer system. Membranes were blocked with 5% skim milk in TBST for 1 h at room temperature and incubated with a single primary antibody diluted in 5% BSA overnight at 4°C. Primary antibodies included anti–TDP-43 (Abnova, #H00023435-M01, 1:1000) and anti-GAPDH (Cell Signaling Technology, #2118, 1:1000). The following day, membranes were washed with 1× TBST and incubated with secondary antibody (Cytiva, #NA931, 1:10,000) diluted in blocking buffer for 1 h at room temperature. Protein signals were detected using ECL reagents (Vazyme, #E423; Bio-Rad, #1705061) and imaged with a Bio-Rad ChemiDoc system.

### Basescope assay

BaseScope in situ hybridization assay was performed using the BaseScope v2 Detection Reagent Kit–RED (Advanced Cell Diagnostics, #323900) according to the manufacturer’s instructions with minor modifications. Custom 1ZZ probes were designed to specifically target the back-splice junction sequences of circADGRL3 and circZNF566.

Formalin-fixed, paraffin-embedded (FFPE) human postmortem brain sections were obtained from the Johns Hopkins Brain Resource Center. Slides were deparaffinizated in xylene and rehydration through a graded ethanol series. Sections were treated with hydrogen peroxide for 12 min at room temperature and subjected to target retrieval in RNAscope Target Retrieval Reagent at ∼99 °C for 15 min. After washing, samples were incubated with RNAscope Protease III at room temperature for 30 min to permeabilize tissue. Positive and negative control probes (provided in the kit), along with target-specific probes, were hybridized to sections in parallel for 2 h at 40 °C in a humidified oven. Signal amplification was carried out through sequential amplification steps (AMP1–AMP8), followed by detection using Fast RED chromogenic substrate. Slides were counterstained with hematoxylin, air-dried, and mounted using VectaMount (Vector Laboratories, #H-5000).

Images were acquired using a brightfield microscope at 60× magnification. RNA signals were visualized as discrete punctate red foci, each representing a single RNA molecule. Signal specificity was validated using positive control probes and negative control probes.

For quantification, at least 10 fields of view were imaged per section. The total number of cells was determined based on nuclear counterstaining. Cells with small, hyperchromatic nuclei, characteristic of glial morphology, were excluded based on nuclear size and staining intensity. A minimum of 100 cells were quantified per section. The number of red foci was counted and summed for each section, and normalized to total cell number. All image acquisition and quantification were performed in a blinded manner, with the operator unaware of disease condition.

### Statistical analysis

Quantification methods, explaining of n and type of statistical test is indicated in each figure legend. Analyzed data was plotted and tested for statistical significance using the GraphPad Prism software and RStudio. P value of < 0.05 was considered to be significant (*P < 0.05, ** P < 0.01, *** P < 0.001, **** P < 0.0001).

