## Supplemental information for "Loss of TDP-43 function drives cryptic circular RNAs in neurodegenerative diseases"

### i. Supplementary Figures

**Figure S1.** Validation of representative cryptic circRNAs regulated by TDP-43 depletion.

**Figure S2.** Identification of TDP-43 binding sites by ARTR-seq and genomic features associated with cryptic circRNA formation.

**Figure S3**. Experimental design for time-course RNA analysis and validation of subcellular fractionation in $i^3$Neurons.

**Figure S4.** Expression of host mRNAs corresponding to representative cryptic circRNAs in human postmortem brain tissues.

### ii. Supplementary Tables

**Table S3.** Validated full sequences of selected cryptic circRNAs (5'-3').

**Table S5**. Demographic information for formalin-fixed paraffin embedded (FFPE) and frozen postmortem tissues.

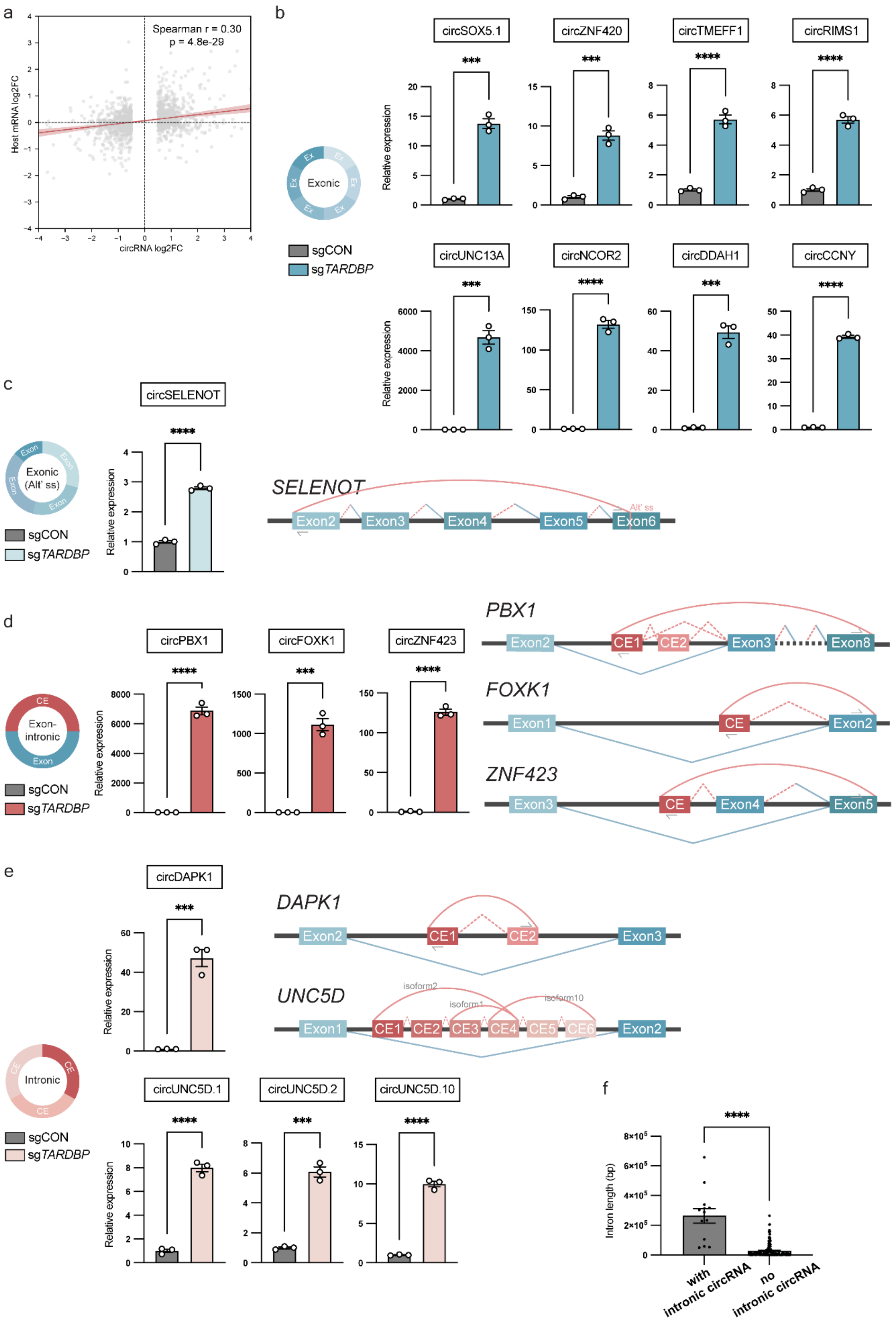
a
Spearman r = 0.30
p = 4.8e-29
Host mRNA log2FC
circRNA log2FC
b
circSOX5.1
circZNF420
circTMEFF1
circRIMS1
circUNC13A
circNCOR2
circDDAH1
circCCNY
Relative expression
Exonic
sgCON
sgTARDBP
c
circSELENOT
Exonic (Alt' ss)
SELENOT
Alt' ss
Exon2
Exon3
Exon4
Exon5
Exon6
d
circPBX1
circFOXK1
circZNF423
Exon-intronic
PBX1
CE1
CE2
Exon3
Exon8
FOXK1
Exon1
CE
Exon2
ZNF423
Exon4
Exon5
e
circDAPK1
DAPK1
UNC5D
isoform2
isoform1
isoform10
CE3
CE4
CE5
CE6
Intronic
circUNC5D.1
circUNC5D.2
circUNC5D.10
f
Intron length (bp)
with intronic circRNA
no intronic circRNA

**Figure S1. Validation of representative circRNAs regulated by TDP-43 depletion.**

**a,** Correlation between the abundance of significantly changed circRNAs and expression of their corresponding host mRNAs following TDP-43 depletion in $i^3$Neurons. Each dot represents an individual gene. CircRNA (the most abundant isoform for each gene) and host gene expression changes are shown as log2 fold change. The red line indicates linear regression with shaded area representing the 95% confidence interval. Spearman correlation coefficient and associated P value are shown.

**b–e,** Splicing schemes and RT–qPCR validations of additional upregulated circRNAs representing distinct categories. Rounded arcs indicate back-splicing events, whereas angular arcs indicate linear splicing events. Blue arcs denote canonical splice junctions, red arcs denote cryptic circRNA-associated junctions, and bi-colored blue–red arcs indicate splice junctions shared between canonical and circular transcripts. Solid arcs represent experimentally validated or canonical splice junctions, whereas dashed arcs indicate predicted splice junctions. Arrows indicate divergent primer positions used for RT–qPCR analyses. Data are shown as mean ± SEM from three independent biological replicates. Statistical significance was determined using two-tailed unpaired t-tests. *P < 0.05, ** P < 0.01, *** P < 0.001, **** P < 0.0001.

**b,** Validation of exonic circRNAs including circSOX5.1, circZNF420, circTMEFF1, circRIMS1, circUNC13A, circNCOR2, circDDAH1, and circCCNY.

**c,** Validation of circSELENOT, an alternatively spliced exonic circRNA generated through activation of an alternative splice site within an annotated exon. The alternative splice site is indicated by a horizontal red line.

**d,** Validation of exon–intronic cryptic circRNAs containing cryptic exon sequences and canonical exons, including circPBX1, circFOXK1, and circZNF423.

**e,** Validation of intronic cryptic circRNAs derived predominantly from cryptic exon-like sequences within introns, including circDAPK1 and multiple circUNC5D isoforms. Distinct circRNA isoforms generated from alternative cryptic splice-site combinations are indicated.

**f,** Comparison of intron lengths associated with intronic cryptic circRNAs and other introns within the same host genes. Introns giving rise to intronic cryptic circRNAs were significantly longer than other introns from the corresponding genes. Data are shown as individual introns with bars indicating mean ± SEM. Statistical significance was determined using a two-tailed Mann–Whitney test. ****$P < 0.0001$.

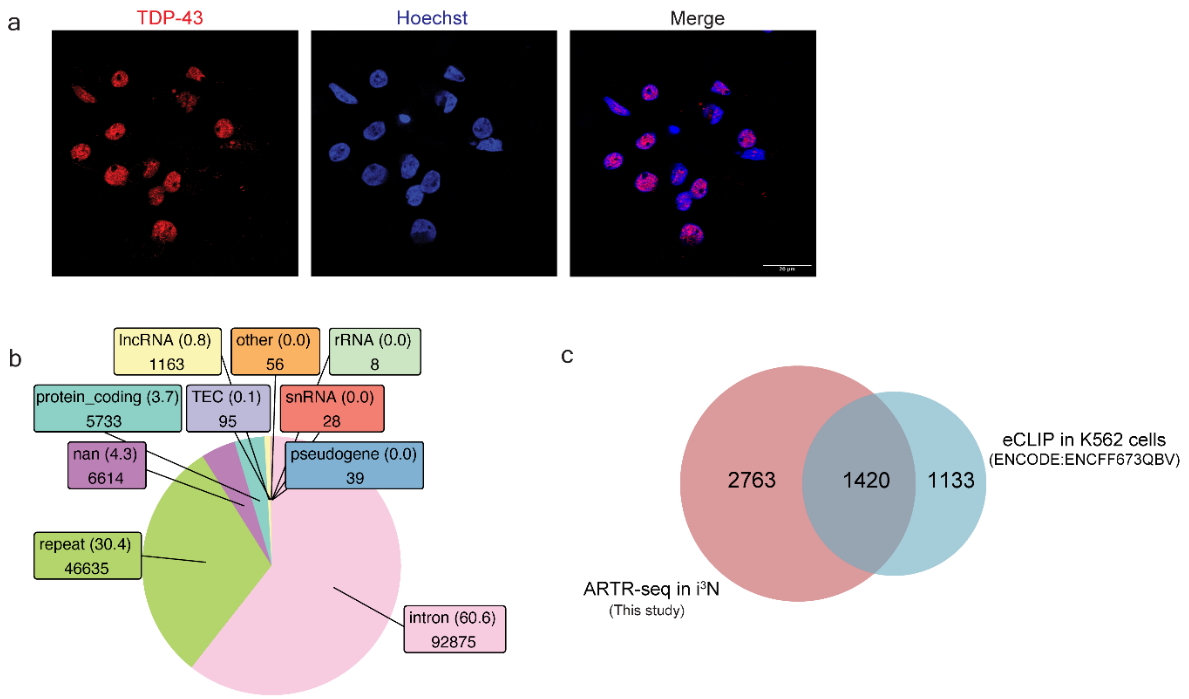


**Figure S2. Identification of TDP-43 binding sites by ARTR-seq and genomic features associated with cryptic circRNA formation.**

**a,** Representative immunofluorescence images showing TDP-43 staining used for ARTR-seq in i$^3$Neurons. TDP-43 is shown in red and nuclei are counterstained with Hoechst (blue). Scale bar, 10 µm.

**b,** Genomic annotation of ARTR-seq peaks identified in control i$^3$Neurons. The majority of TDP-43–associated regions mapped to intronic sequences, followed by repetitive elements and unannotated regions. Numbers indicate peak counts and their percentages of all identified peaks.

**c,** Overlap between TDP-43–associated genes identified by ARTR-seq in i$^3$Neurons and previously published TDP-43 eCLIP data from K562 cells (ENCODE dataset ENCFF673QBV). The substantial overlap supports the specificity of ARTR-seq–identified TDP-43 binding sites.

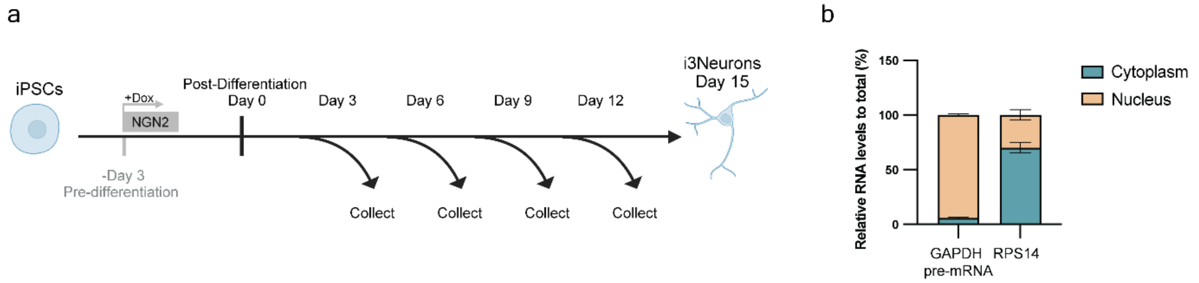


**Figure S3. Experimental design for time-course RNA analysis and validation of subcellular fractionation in i$^{3}$Neurons.**

**a,** Schematic of the i$^{3}$Neuron differentiation protocol used for time-course RNA analysis. Human iPSCs were induced with doxycycline-inducible NGN2 beginning 3 days before differentiation (Day -3). Day 0 denotes the start of neuronal differentiation. Cells were collected on Days 3, 6, 9, 12, and 15 for downstream analyses.

**b,** Validation of nuclear and cytoplasmic fractionation in TDP-43–depleted i$^{3}$Neurons. Relative abundance of the nuclear-enriched transcript GAPDH pre-mRNA and the cytoplasmic-enriched ribosomal transcript RPS14 was measured by RT–qPCR normalized to the RNA spike-in control and presented relative to total RNA. Data are presented as percentage of total RNA recovered from both fractions and shown as mean ± SEM from four biological replicates.

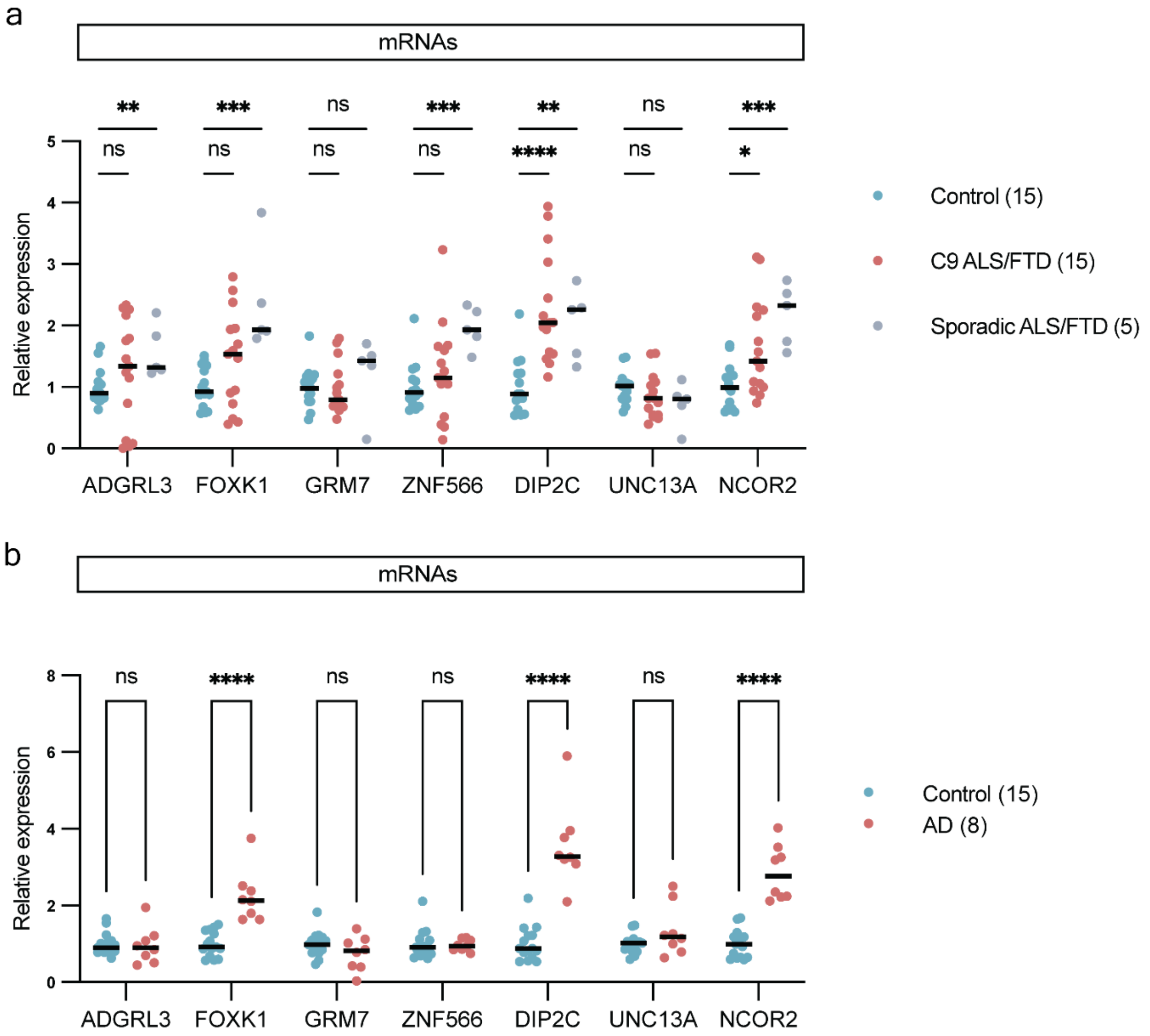


**Figure S4. Expression of host mRNAs corresponding to representative cryptic circRNAs in human postmortem brain tissues.**

**a,** RT–qPCR analysis of host mRNA expression in temporal cortex tissues from non-neurological controls (n = 15), C9ORF72-associated ALS/FTD (n = 15), and sporadic FTD cases (n = 5). Relative expression levels of *ADGRL3*, *FOXK1*, *GRM7*, *ZNF566*, *DIP2C*, *UNC13A*, and *NCOR2* mRNAs were normalized to HPRT1. Most host mRNAs showed no significant changes or modest alterations between disease groups and controls despite robust upregulation of the corresponding cryptic circRNAs. Each dot represents a sample from an individual donor. Horizontal bars indicate median values. Statistical significance was determined using two-tailed Mann–Whitney tests. ns, not significant. *$P < 0.05$, **** $P < 0.0001$.

**b,** RT–qPCR analysis of host mRNA expression in temporal cortex tissues from non-neurological controls (n = 15) and Alzheimer's disease (AD) cases (n = 8). Relative expression levels of mRNAs were normalized to HPRT1. *FOXK1*, *DIP2C*, and *NCOR2* mRNAs were modestly elevated in AD tissues, whereas *ADGRL3*, *GRM7*, *ZNF566*, and *UNC13A* mRNAs were not significantly altered. Each dot represents a sample from an individual donor. Horizontal bars indicate median values. Statistical significance was determined using two-tailed Mann–Whitney tests. ns, not significant. **** $P < 0.0001$.

**Table S3, Validated full sequences of selected cryptic circRNAs (5'-3').** Uppercase letters indicate exonic sequences, whereas lowercase letters indicate intronic cryptic sequences.

**circADGRL3.2**

auuuaugucuucauuucuugcauuugaaguacucuucgaugacauccuuggccugagacuccuugccauaguccuuaacuacugcacaacugcaaccaaccacuuuaugaggcuuccccucucugucaguuuuacagaggccuacccuuuccccuaguuucuuguugucaucaaccuuaauuagguugauuugguguucagcacaaagggccuccaccaacuugacauacaucacaggcucaucacaguuggaugcaagcacacaaaggugggcuuggugcuugucuaagccuuugguagcuucaugaauuccacgugcuaggcaaucguggaugagggcagucuucagcaccuccuguaaagcaguauuaacguccauuacgcuuccagcagcaauaccuuccucggccauggcgguggguuaccggugaagacgaaucuuguuuuuauuuauuuauuuauuuauuuauuuauuuuuuugagacggagucucacucugucgcccaggcuggagugcaguggcacgaucucggcucacugcaagcuccgccucgaggguucaugccauucuccugucucggccucccgaguagcugggacuccaggcgcccgccaccacacccggcuaauuuuuuguauuuuuaguagagacggg guuucacuguguuagccaggaugaucucgaucuccugaccuugugauccacccgccucggccucccaaagugcugggauuacaggcgugagccacggcgccaagcccugaagccgaaucuuaaacgcacccaagccuccgccucugcacgacucggcggcagcagggaaagaggccucacgcgccgaaaagGGCAGGCACAUCAUGGACAAGUUUCAUACAUUUCUCCGCCAAUUCACCUUGACUCUGAGCUAGAAAGACCCUCUGUUAAAG

**circGRM7.1**

aucugugguuacaacucuaucuacagauauauuugcugaagaaaauauuucuuccaccuuccacagcaaaauuccauucauuuaagaccaauacagcaugucccaaaugucuuaguccaguuuuaaucuuuaauagcuucagaagugugaaauuacaaacaugccccccaaaaaucauuugaaaguuuaauuauuuaaauuucucuuuguauuuacauuuguggauuuugaaaaaucaauauuuaauuguacuuuguuuguugcuguccuucugauugaagauggcaaacauugacuuaaagaccaaaaaguuguaacuaaauauguauuauucaaaauuaucaaaacuaaauaagugaccaagaaauucaaaugauuagacuuucaaaggugauuuuuuauguguuugcgugugugugugugugugugugugugugugugugugugugugguaguauuuuuucuucugaagaaacugaucuugaaacucuacuaagauuuuagggacaucuaguuuaaugagcaaaugcuuugugucugggacugcagcaugcacuagauauuuaagagauuaacau aaaauaauaauaauaauacaaaaacaacaacacauccucucagacauggaagaaaauuccacuugagaaauggaccaauacaucacgcacuaauaaaauagcaaaggcucucaaacaucaaugugcugaagaaucacauggcaagcaugccaagcagauguccuggcgggcuuuagaaacuucuggaagaaguggacagagaauucuuaugguucacauauaaacaauaugaaucuuuggaagaaaaauucauuuagggaccaaaagccccuuuauaucugauuuuuuucccucucaccaaauuuucacaaauuuuccuuuccaaaaugccuuugagcaaaucuguuugauaucaccccauccacuucuccugucugaauucuuuauuacuaagaauccuucagcauuccaucucucacucgucuuugucaaaagcauuucucagugucuuuauuccuucucgucucguauuuggacucaucucucaaacugaaaaaggagauggcuaugguuucuccagaagcuuggagaacuucccuuugcaaaaaaagguagcgauguuaauuaacuugaaagguagaacuagaugacauugcaggcagugagauguuaucauguuugaccccuggaacuguccaaaaauggaagaagcugccucaucag

**circZNF566.2**

CUCUACCCUUCUCCAAAAGAAGAGCCAAGAGAAGGUCCUUUUCUACAAAUAUCAGAGCCAUGGCUCAGGAGUCAGUGAUGUUCAGUGAUGUGUCCGUAGACUUCUCUCAGGAGGAGUGGGAAUGCCUGAAUGAUGAUCAGAGAGAUUUAUACAGAGAUGUGAUGUUGGAGAAUUACAGCAACCUGGUUUCAAUGGCAGGGCAUUCUAUUUCUAAACCAAAUGUGAUCUCCUACUUGGAGCAAGGGAAGGAGCCCUGGUUGGCUGACAGAGAGCUAACAAGAGGCCAGUGGCCAGUCCUGGAAUCAAGAUGUGAGACCAAGAAAUUAUUUCUGAAGAAAGAAAUUUAUGAAAUAGAAUCAACCCAGUGGGAAAUAAUGGAAAAACUCACAAGACGUGAUUUUCAGUGCUCCAGUUUCAGAGAUGAUUGGGAAUGUAAUCGGCAGUUUAAGAAAGAACUCGGCUCUCAGGGGGGACAUUUCAAUCAAUUGGUAUUCACUCAUGAAGAUCUGCCCACUUUGAGUCACCAUCCAUCCUUCACAUUACAGCAAAUCAUUAACAGUAAAAAGAAAUUCUGUGCAUCUAAAGAAUAUAGGAAAACCUUUAGACAUGGCUCACAGUUUGCUACACAUGAGAUAAUUCAUACCAUUGAGAAGCCUUAUGAAUGUAAGGAAUGUGGAAAGUCCUUUAGACAUCCCUCAAGACUC

ACUCAUCAUCAGAAAAUUCAUACUGGCAAGAAACCCUUUGAAUGUAAGGAAUGUGGAAAAACCUUUAUUUGUGGCUCAGACCUUACUCGACAUCACAGAAUUCACACUGGUGAGAAACCCUAUGAAUGUAAGGAAUGUGGGAAAGCCUUUAGUAGUGGUUCAAACUUCACUCGACAUCAGAGAAUUCACACAG

**circDIP2C.3**

GUGAGUCGCUCUGCCUGUCUGAUGACGACACAGCUGAUCUGUAAGUUGCUGCGGUCCAGGGAGGCGGCGGCUGUGGACGUCAGGACGUGGCCCCUCAUCCUGGACACAGAUGAUUUGCCAAAGAAGCGGCCUGCCCAGAUCUGCAAACCUUGCAACCCAGACACUCUUGCAUAUCUCGACUUCAGCGUGUCCACAACUGGGAUGCUAGCUGGCGUAAAGAUGUCUCACGCAGCCACCAGUGCCUUCUGCCGUUCCAUUAAGCUGCAGUGUGAACUUUACCCCUCUAGAGAAGUGGCCAUCUGCCUGGACCCUUACUGUGGACUGGGAUUUGUCCUCUGGUGCCUCUGCAGUGUGUAUUCUGGGCACCAGUCCAUCCUGAUCCCGCCCUCUGAGCUGGAAACCAACCCCGCCUUGUGGCUUCUUGCCGUGAGUCAGUACAAAGUCCGAGACACGUUUUGCUCCUACUCCGUGAUGGAGCUGUGCACCAAGGGGCUGGGCUCGCAAACAGAGUCCCUCAAGGCGCGAGGGCUGGACUUGUCCCGAGUGAGGACCUGCGUGGUUGUGGCGGAAGAGAGGCCUCGGAUCGCACUCACACAGUCGUUCUCAAAGCUGUUUAAGGACCUGGGCCUUCACCCGCGGGCCGUCAGCACCUCGUUCGGUUGCAGGGUGAACCUGGCGAUUUGCUUGCAGGGAACCUCAGGACCUGACCCAACCACUGUCUACGUGGACAUGAGAGCCCUGAGACACGACAG

**Table S5, Demographic information for formalin-fixed paraffin embedded (FFPE) and frozen postmortem tissues.**

| Tissue ID | Patient GUID | Clinical Diagnosis | Age at time of death | Gender | Experiment usage |
|---|---|---|---|---|---|
| Control-1 | BRC1613 | Non-neurologic control | 74 | Male | Frozen |
| Control-2 | BRC2052 | Non-neurologic control | 79 | Male | Frozen, FFPE |
| Control-3 | BRC2103 | Non-neurologic control | 88 | Male | FFPE |
| Control-4 | BRC2143 | Non-neurologic control | 86 | Male | FFPE |
| Control-5 | BRC2151 | Non-neurologic control | 72 | Male | Frozen |
| Control-6 | BRC2209 | Non-neurologic control | 71 | Male | Frozen |
| Control-7 | BRC2228 | Non-neurologic control | 64 | Male | Frozen |
| Control-8 | BRC2234 | Non-neurologic control | 68 | Female | Frozen |
| Control-9 | BRC2317 | Non-neurologic control | 65 | Female | Frozen |
| Control-10 | BRC1058 | Non-neurologic control | 70 | Male | Frozen |
| Control-11 | BRC1273 | Non-neurologic control | 68 | Male | Frozen |
| Control-12 | JHU100 | Non-neurologic control | 52 | Female | Frozen |
| Control-13 | JHU101 | Non-neurologic control | 70 | Female | Frozen |
| Control-14 | JHU110 | Non-neurologic control | 50 | Male | Frozen |
| Control-15 | JHU123 | Non-neurologic control | 52 | Male | Frozen |
| Control-16 | JHU124 | Non-neurologic control | 85 | Male | Frozen |
| Control-17 | JHU129 | Non-neurologic control | 63 | Female | Frozen |
| C9-FTD/ALS-1 | BRC1814 | MND/Dementia | 64 | Male | Frozen |

| | | | | | |
|---|---|---|---|---|---|
| C9-FTD/ALS-2 | BRC1909 | FTD-MND | 71 | Female | Frozen |
| C9-FTD/ALS-3 | BRC2500 | FTLD-MND | 63 | Female | FFPE |
| C9-FTD/ALS-4 | BRC2589 | FTLD-MND/MNI | 72 | Female | Frozen |
| C9-FTD/ALS-5 | BRC2655 | FTLD | 79 | Male | Frozen, FFPE |
| C9-FTD/ALS-6 | BRC2665 | FTLD | 62 | Male | Frozen |
| C9-FTD/ALS-7 | BRC2696 | FTD | 64 | Female | Frozen, FFPE |
| C9-FTD/ALS-8 | BRC2770 | FTLD-MND/MNI | 67 | Male | Frozen |
| C9-FTD/ALS-9 | BRC2713 | FTLD-MND/MNI | 70 | Male | Frozen |
| C9-FTD/ALS-10 | BRC2735 | FTD | 58 | Female | Frozen |
| C9-FTD/ALS-11 | JHU88 | FTLD/fALS | 59 | Male | Frozen |
| C9-FTD/ALS-12 | JHU119 | FTD/fALS | 61 | Female | Frozen |
| C9-FTD/ALS-13 | JHU19 | fALS | 52 | Male | Frozen |
| C9-FTD/ALS-14 | JHU22 | fALS | 66 | Male | Frozen |
| C9-FTD/ALS-15 | JHU92 | fALS | 72 | Male | Frozen |
| C9-FTD/ALS-16 | JHU120 | fALS | 68 | Female | Frozen |
| Sporadic FTD-1 | BRC2392 | Sporadic FTD-TDP | 78 | Male | Frozen |
| Sporadic FTD-2 | BRC2621 | Sporadic FTD-TDP | 64 | Female | Frozen |
| Sporadic FTD-3 | BRC2640 | Sporadic FTD-TDP | 84 | Male | Frozen |
| Sporadic FTD-4 | BRC2667 | Sporadic FTD-TDP | 74 | Male | Frozen |
| Sporadic FTD-5 | BRC2802 | Sporadic FTD-TDP, mixed (AD+PD) | 78 | Female | Frozen |
| AD-1 | BRC2277 | Alzheimer's disease | 63 | Female | Frozen |
| AD-2 | BRC2282 | Alzheimer's disease | 79 | Female | Frozen |

| | | | | | |
|---|---|---|---|---|---|
| AD-3 | BRC2299 | Alzheimer’s disease | 79 | Male | Frozen |
| AD-4 | BRC2365 | Alzheimer’s disease | 79 | Male | Frozen |
| AD-5 | BRC2376 | Alzheimer’s disease | 61 | Male | Frozen |
| AD-6 | BRC2417 | Alzheimer’s disease | 61 | Female | Frozen |
| AD-7 | BRC2419 | Alzheimer’s disease | 79 | Male | Frozen |
| AD-8 | BRC2609 | Alzheimer’s disease | 68 | Male | Frozen |
